# vcfgl: A flexible genotype likelihood simulator for VCF/BCF files

**DOI:** 10.1101/2024.04.09.586324

**Authors:** Isin Altinkaya, Rasmus Nielsen, Thorfinn Sand Korneliussen

## Abstract

**Motivation:** Accurate quantification of genotype uncertainty is pivotal in ensuring the reliability of genetic inferences drawn from NGS data. Genotype uncertainty is typically modeled using Genotype Likelihoods (GLs), which can help propagate measures of statistical uncertainty in base calls to downstream analyses. However, the effects of errors and biases in the estimation of GLs, introduced by biases in the original base call quality scores or the discretization of quality scores, as well as the choice of the GL model, remain under-explored.

**Results:** We present vcfgl, a versatile tool for simulating genotype likelihoods associated with simulated read data. It offers a framework for researchers to simulate and investigate the uncertainties and biases associated with the quantification of uncertainty, thereby facilitating a deeper understanding of their impacts on downstream analytical methods. Through simulations, we demonstrate the utility of vcfgl in benchmarking GL-based methods. The program can calculate GLs using various widely used genotype likelihood models and can simulate the errors in quality scores using a Beta distribution. It is compatible with modern simulators such as msprime and SLiM, and can output data in pileup, VCF/BCF and gVCF file formats. The vcfgl program is freely available as an efficient and user-friendly software written in C/C++.

**Availability:** vcfgl is freely available at https://github.com/isinaltinkaya/vcfgl.

**Supplementary information:** Supplementary information is available online.

## 1 Introduction

Next-generation sequencing (NGS) has enabled a deeper understanding of genetic variation across time and various biological systems, particularly non-model organisms and ancient populations (da Fonseca *et al*., 2016). Accurately quantifying genotype uncertainty through genotype likelihoods is fundamental in ensuring the reliability of genetic inferences drawn from NGS data. This is particularly important in low-depth and ancient DNA data analysis, where the biases and genotype uncertainties are pronounced, and the probability of only sampling nucleotides from one chromosome in a diploid is non-negligible.

Genotype likelihoods provide a probabilistic measure of genotype uncertainty that incorporates information regarding alignment or assembly uncertainty and base-calling uncertainty (Nielsen *et al*., 2011). Genotype likelihoods are integral not only to calling genotypes but also to a multitude of downstream methods used in a diverse set of scientific inquiries, such as estimating relatedness (Korneliussen and Moltke, 2015; Waples *et al*., 2019), allele frequency spectra (Korneliussen *et al*., 2014; Mas-Sandoval *et al*., 2022; Rasmussen *et al*., 2022), calculating genetic distances (Vieira *et al*., 2016; Zhao *et al*., 2022), conducting Principal Component Analysis (Meisner and Albrechtsen, 2018), evaluating linkage disequilibrium (Fox *et al*., 2019), and admixture proportions (Skotte *et al*., 2013; Lou *et al*., 2021; O’Rawe *et al*., 2015).

The process of quantifying uncertainty using quality scores and genotype likelihoods is in itself subject to uncertainties and potential biases caused by errors or biases in the estimation of per base error probabilities or arising from the discretization of genotype likelihood values and choice of genotype likelihood model. Such factors introduce additional layers of bias and uncertainty, which may not have been considered or be sufficiently addressed by existing methods and frameworks. Moreover, new sequencing platforms and continuous changes in sequencing technologies necessitate flexible tools for quantifying the effect of estimation uncertainty through simulations.

Existing tools for simulating genotype likelihoods, such as msToGlf (utility program in ANGSD package), have contributed to developing and evaluating various genotype likelihood-based methods (Zhao *et al*., 2023; Soraggi *et al*., 2018; Luqman *et al*., 2021; Wang *et al*., 2016; Mas-Sandoval *et al*., 2022; Fox *et al*., 2019; Korneliussen *et al*., 2014). However, they do not enable modeling of uncertainty in the estimation of genotype likelihoods. Moreover, they are often not designed to be compatible with state-of-art simulation tools and modern file types such as VCF/BCF files. This gap in the simulation capabilities limits researchers’ ability to effectively examine and benchmark the effects of the quantification of uncertainty in NGS data, underscoring the need for modern and flexible tools to model these uncertainties and simulate data.

## 2 Method

We introduce vcfgl, a lightweight utility tool for simulating genotype likelihoods. The program incorporates a comprehensive framework for simulating uncertainties and biases, including those specific to modern sequencing platforms. It offers compatibility with modern simulators such as msprime (Baumdicker *et al*., 2022) and SLiM (Messer, 2013; Haller and Messer, 2023) through the use of VCF files. It is a lightweight tool that does not require many dependencies and is implemented in C/C++ for facilitating fast and efficient simulations. To our knowledge, vcfgl is the only tool for simulating genotype likelihoods offering this functionality. The resulting VCF files can then be used with many tools and frameworks, such as BCFtools (Danecek *et al*., 2021), GATK (McKenna *et al*., 2010; Van der Auwera and O’Connor, 2020), and ANGSD (Korneliussen *et al*., 2014).

Given a VCF file with genotypes, vcfgl can simulate sequencing data, quality scores, calculate the genotype likelihoods and various VCF tags, such as I16 and QS tags used in downstream analyses for quantifying the base calling and genotype uncertainty. For simulating sequence depth, vcfgl uses a Poisson distribution with a fixed mean. For simulating errors in the quality scores, it utilizes a Beta distribution where the shape parameters are adjusted to obtain a distribution with a mean equal to the specified error probability and variance equal to a specified variance parameter (for more details, see Supplementary Material, Section 2). The program provides options for two commonly-used genotype likelihood models, the McKenna genotype likelihood model with independent errors (McKenna *et al*., 2010), and the Li genotype likelihood model that models non-independent error structure and is used in SAMtools/BCFtools (Li, 2011). Detailed descriptions of the models can be found in Supplementary Material, Section 1.

The identification of the variable sites is itself subject to uncertainties, especially in the context of non-model organisms, low-depth sequencing, and ancient DNA data (Nielsen *et al*., 2011). Furthermore, correct handling of invariant and missing sites in downstream analyses is important for the reliability of the conclusions drawn from genomic data. Consequently, the modern and widely utilized Variant Call Format (VCF), originally developed for storing variant information, has evolved to retain the information from invariable sites, thereby facilitating a comprehensive genomic overview. To address this, vcfgl provides the option to simulate the invariable sites, which is usually not possible to obtain directly from simulators. However, as the inclusion of these sites also presents a computational challenge due to the massive increase in the data volume, modern file formats such as genomic VCF (gVCF) have been introduced to address this. The gVCF format uses a genomic block compression approach that can efficiently store invariant sites by grouping them into non-variant block records, thereby reducing the file size footprints (Caetano-Anolles, 2023; Illumina, 2014). Our program can simulate invariable sites, and can output both VCF and gVCF files that are compatible with GATK and BCFtools gVCF formats, thereby allowing the user to both perform analyses incorporating invariable sites and test the effects of various SNP calling methods on downstream analyses.

## 3 Results and discussion

To demonstrate the utility of vcfgl, we benchmarked the accuracy of BCFtools multiallelic genotype calling method under different scenarios mimicking the classic Out-of-Africa model. We simulated variable sites in chromosome 22 for 100 diploid individuals using msprime (Baumdicker *et al*., 2022) with 20 replicates (for more details, see Supplementary Material Section 2). We then used vcfgl to simulate genotype likelihoods and quality scores at read depths of 0.1, 0.5, 1, 2, 10, 20, and 100. We simulated the errors in quality scores using a mean error rate of 0.2% and beta distribution with variance parameters of 0 (no variance, i.e. precise quality scores) and 10^*−*5^. We calculated the associated genotype likelihoods using both Li (`-GL 1`) and McKenna (`-GL 2`) error models. We performed genotype and SNP calling using both naive genotype calling approach, and the BCFtools multiallelic caller (Danecek *et al*., 2021). With the naive genotype caller, we pick the genotype corresponding to the highest genotype likelihood. We used the BCFtools multiallelic genotype caller with the ‘-P 0’ option. The main difference between the two genotype calling methods is then that the BCFtools multiallelic caller identifies alleles (and thereby SNPs) prior to genotype calling and uses the allele frequencies, estimated using the read quality scores, in a Hardy-Weinberg Equilibrium prior for both SNP and genotype calling.(Danecek *et al*., 2016).

To evaluate the performance of genotype calling methods, we calculated two metrics for each simulation replicate for each individual: error rate and call rate. The call rate is calculated as the frequency of the genotype calls for a given threshold for genotype calling, `GQ`. The error rate for a chosen `GQ` is the count of wrongly called genotypes standardized by the count of all genotype calls for the same call criteria. Detailed descriptions regarding the calculation of the error rate and call rate can be found in the Supplementary Material, Section 3.3.

Comparing the two genotype calling approaches, we observe that the additional step of identifying alleles (which includes SNP calling) in the BCFtools multiallelic genotype calling method results in more accurate genotype calls compared to the naive maximum likelihood approach that does not include a SNP calling step. We also observe that with the genotype likelihoods calculated using both McKenna and Li error models, as expected, the error rate decreases with increasing read depth (see Figure 1). At a read depth of 10x, we start to observe higher error rates for the McKenna GL model relative to the Li GL model.

**Fig. 1.**
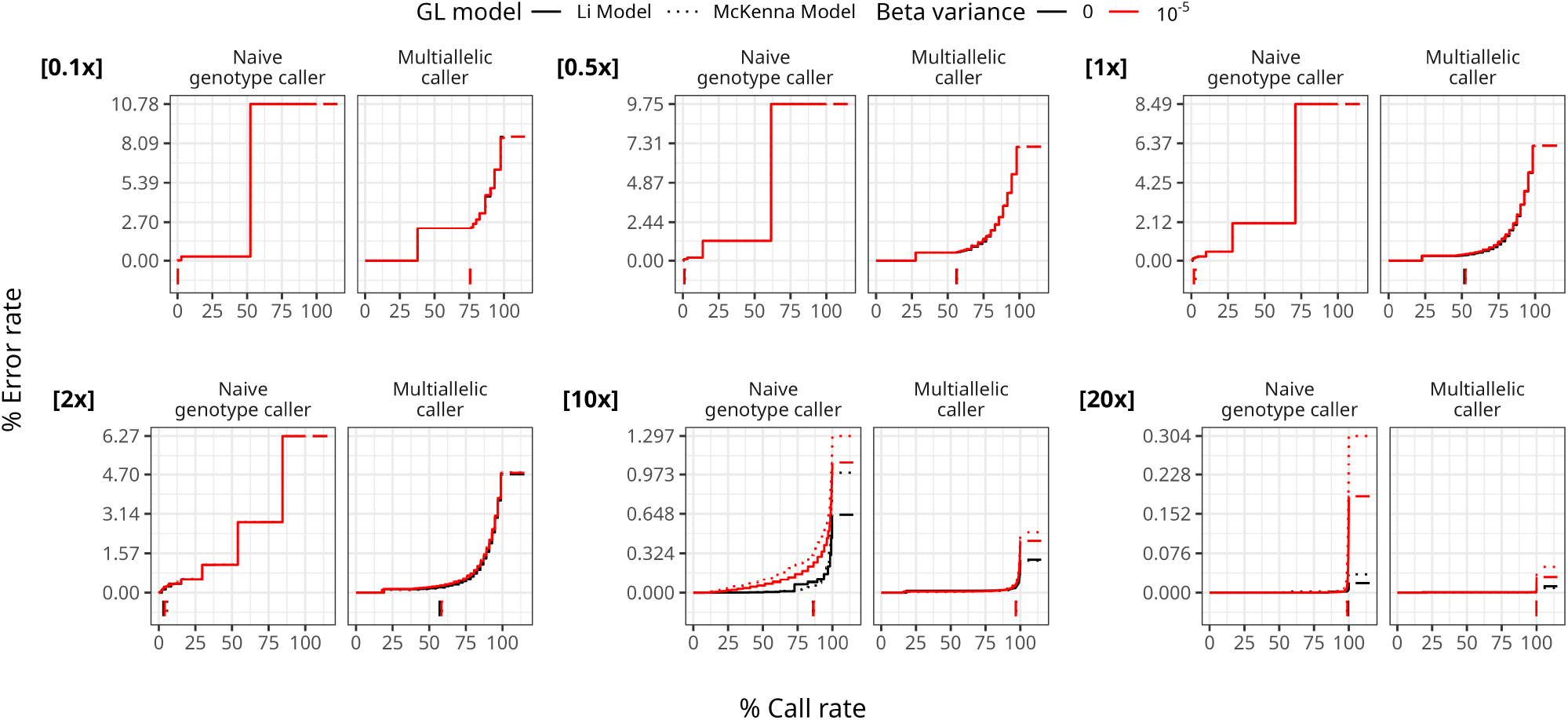
Performance of genotype calling using the naive genotype caller method and BCFtools multiallelic caller with beta distributed errors in the estimation of the quality scores, by sequencing read depth. Colors indicate different variances in the beta distribution (0, and 10^*−*5^, respectively). The line types indicate the Li GL model (1), and McKenna GL model (2) (for details see the Supplementary Material, Section 1.4). The genotype calling error rates (y-axis) and call rates (x-axis) are defined in the main text. The average per-site read depth is indicated in the top left corner of each plot. The curves are obtained by varying the `GQ` threshold for genotype calling. The vertical line segments below 0 on the y-axis denote the minimum `GQ` threshold of 20, and the horizontal line segments after 100 on the x-axis denote the final error rate. The data is from 20 replicates of 100 diploid individuals simulated using msprime, resulting in 328, 230 variable sites per simulation replicate (for details, see the Supplementary Material, Section 2). The BCFtools multiallelic caller is used for each population separately, and the prior parameter is disabled.

Using BCFtools multiallelic caller, we also tested two genotype calling approaches: genotype calling across populations, where the genotype calling is performed on the whole dataset, and within populations genotype calling, where the genotypes are called separately for each population, allowing for population-specific allele frequency estimation and the detection of population-specific variants 1. Across the different read depths, we observe that the within population approach performs better than the across populations approach.

The maximum runtime for these simulations was 7 minutes 40 seconds per replicate without multi-threading, with a mean read depth of 20, when simulating errors in quality scores. Without simulating the quality score errors, the maximum run time was 1 minute 39 seconds. The simulation in both cases consisted of 328, 230 sites and 100 simulated diploid individuals. We addressed the bottleneck in the file writing step by using the HTSlib library’s threading functionality to allow threading in the compression stream. Comparing the use of one thread versus four, we’ve observed a 13% reduction in processing time, down from 7 minutes 40 seconds to 6 minutes 47 seconds. The runtime and file size depend on the number of samples and the amount of sequence data simulated and, of course, on the disk IO. All analyses were conducted on a Red Hat Enterprise Linux 8.8 (Ootpa) system with an Intel(R) Xeon(R) Gold 6152 CPU at 2.10 GHz (x86_64), 754 GiB RAM, and a Linux 4.18.0-477.27.1.el8_8.x86_64 kernel. The benchmarking pipeline was implemented as a Snakemake workflow for reproducibility (Mölder *et al*., 2021) and is freely available.

Future improvements may include modeling alignment and assembly-related biases, for example using mappability maps quantifying mapping biases and site-specific errors.

Our simulation tool, vcfgl, provides a framework for developing more accurate and reliable genetic data analysis methods, ultimately enhancing our understanding of genetic variations and their implications.

---

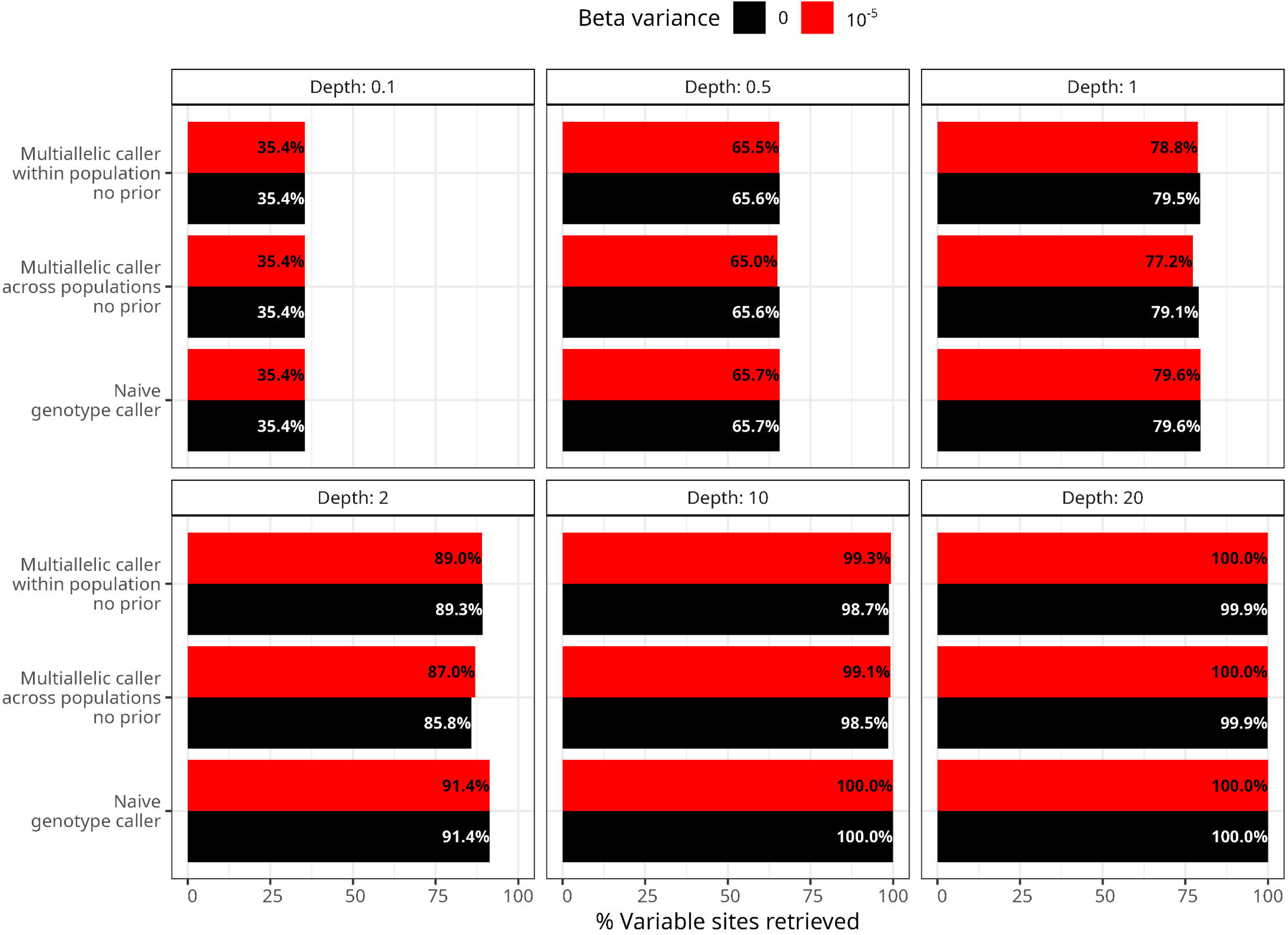

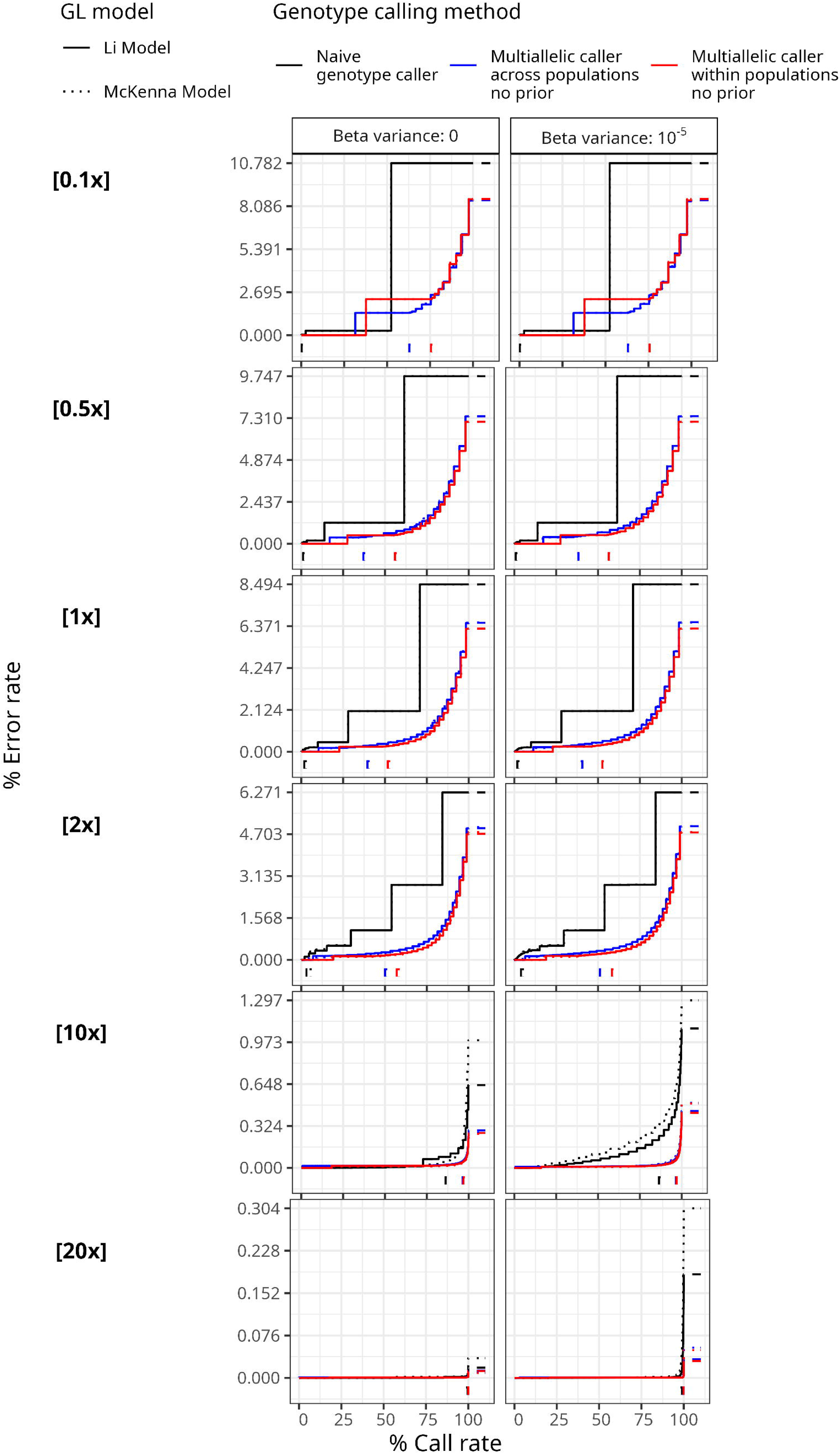

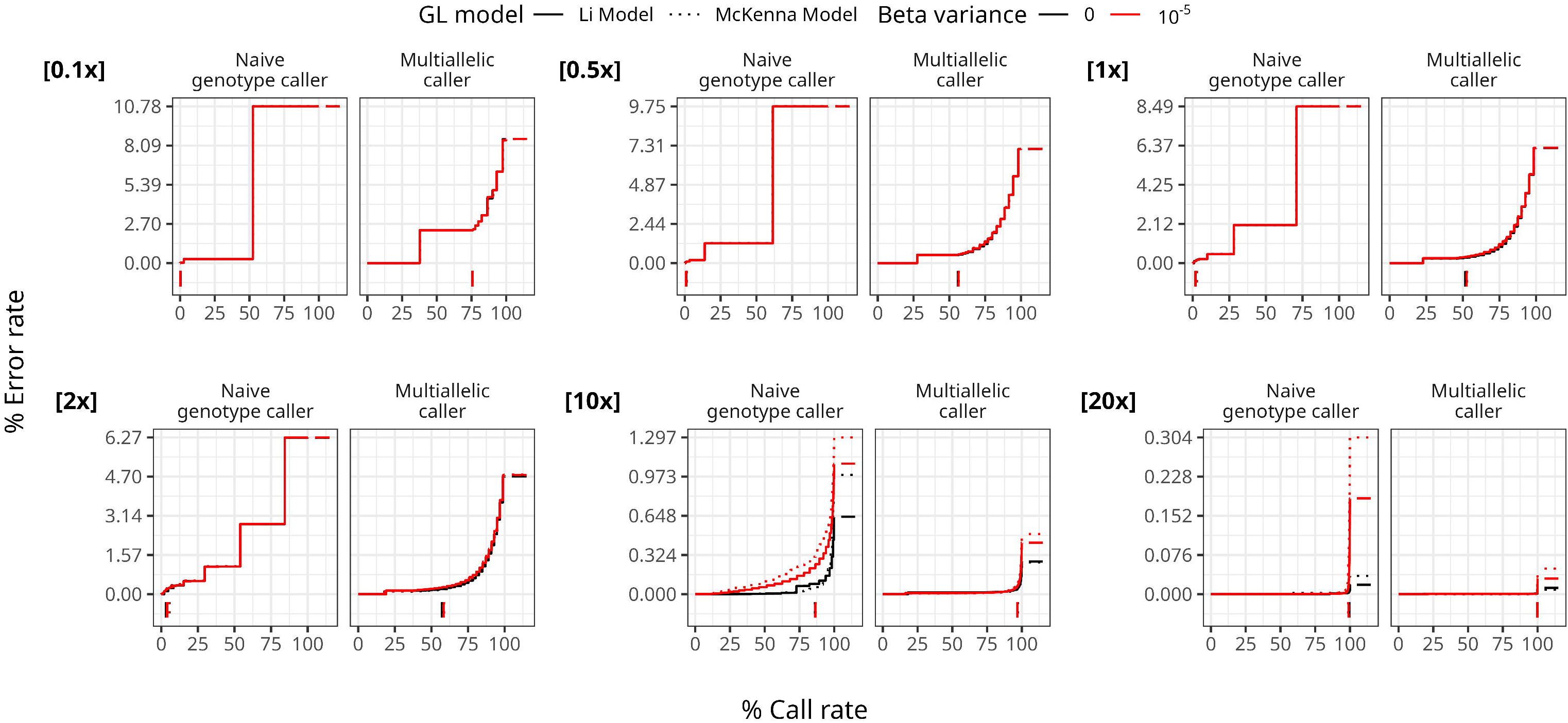

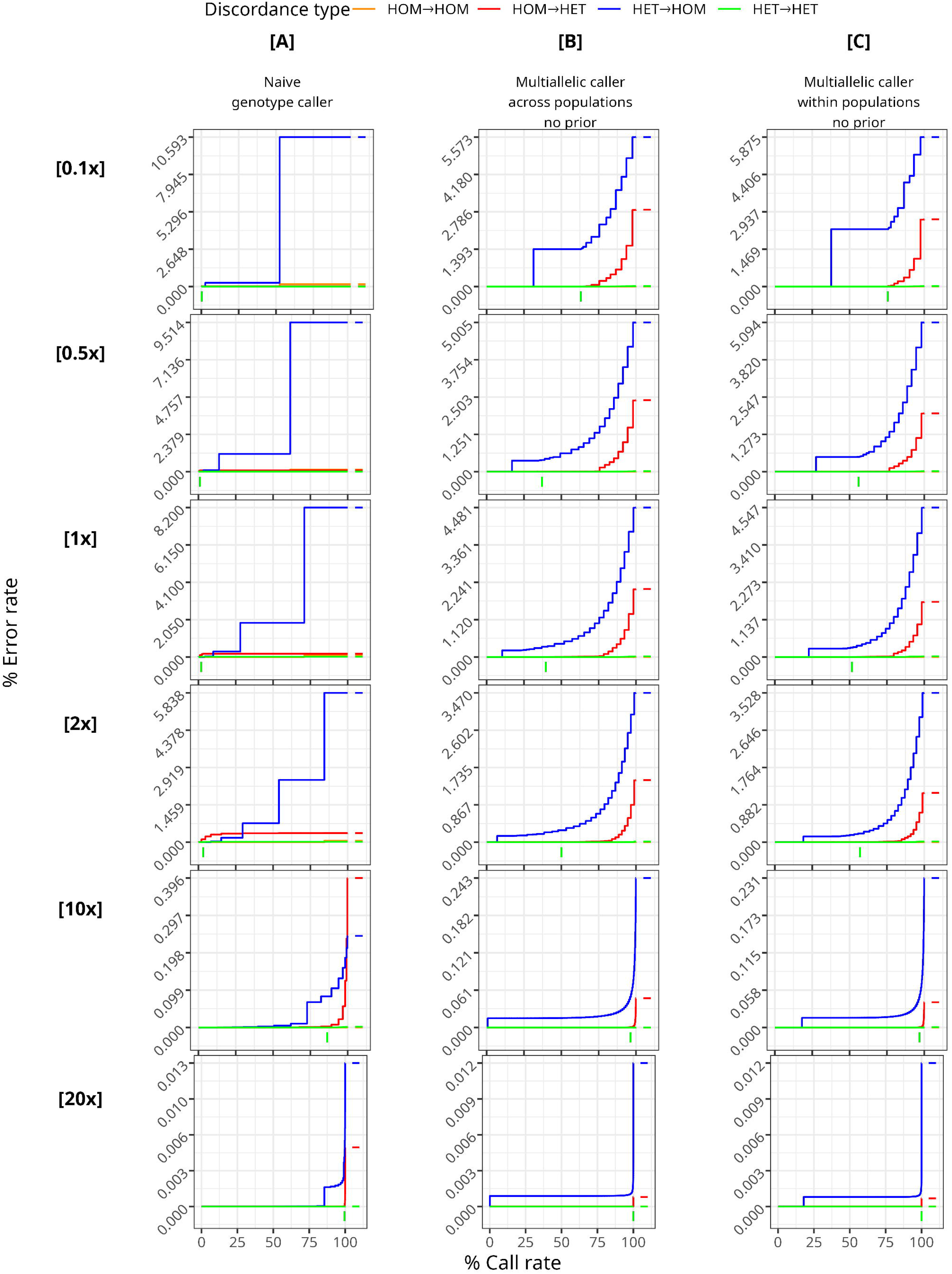

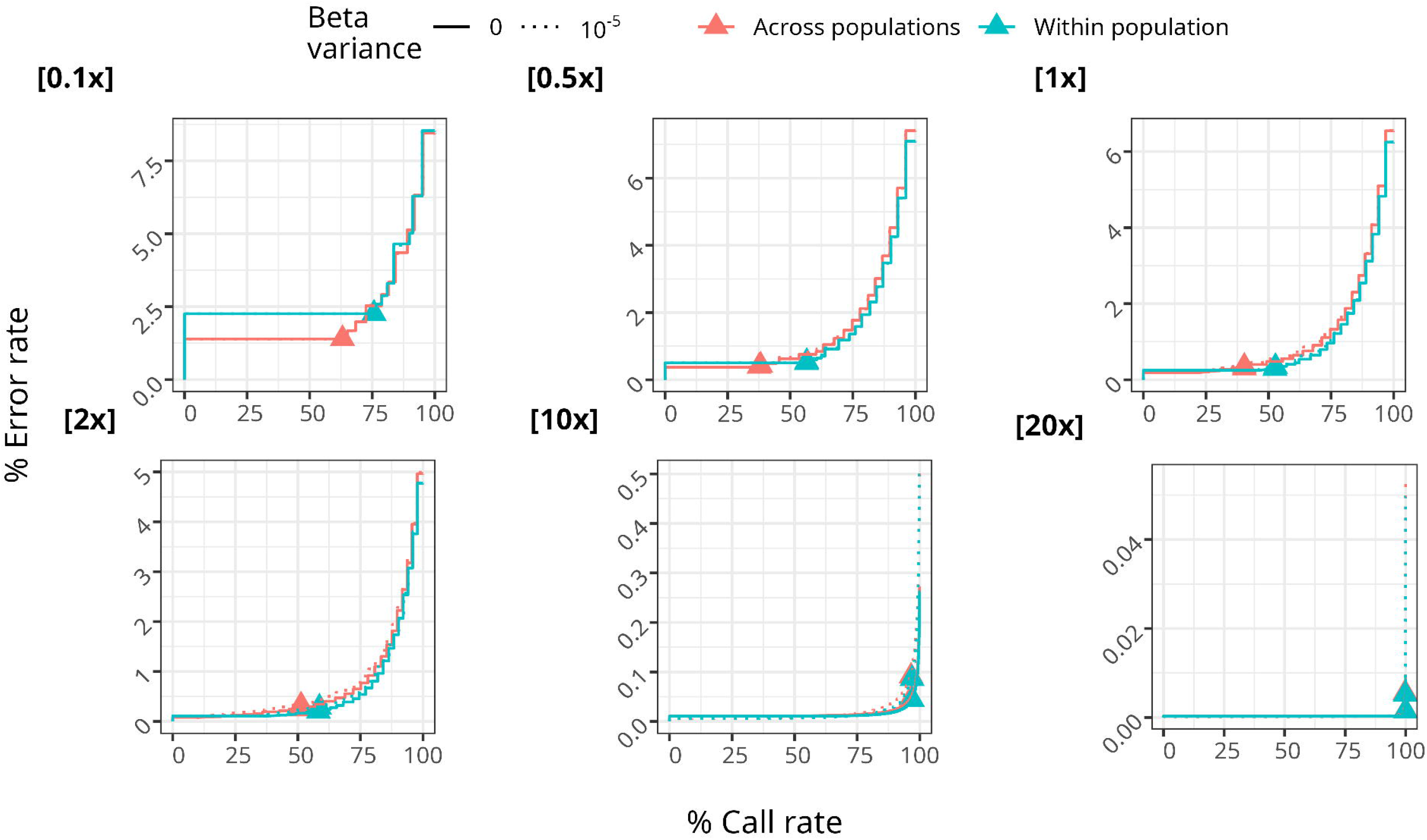

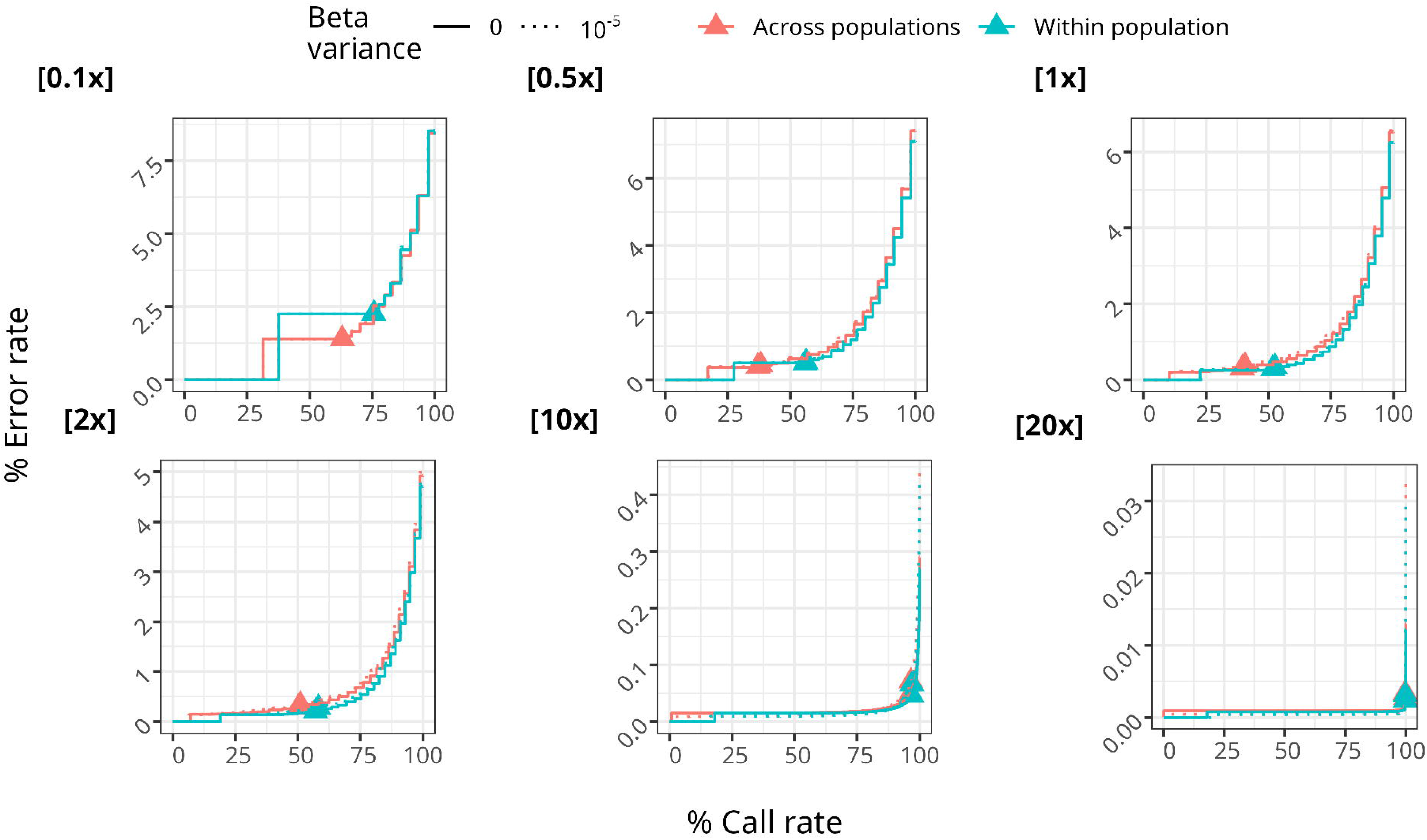

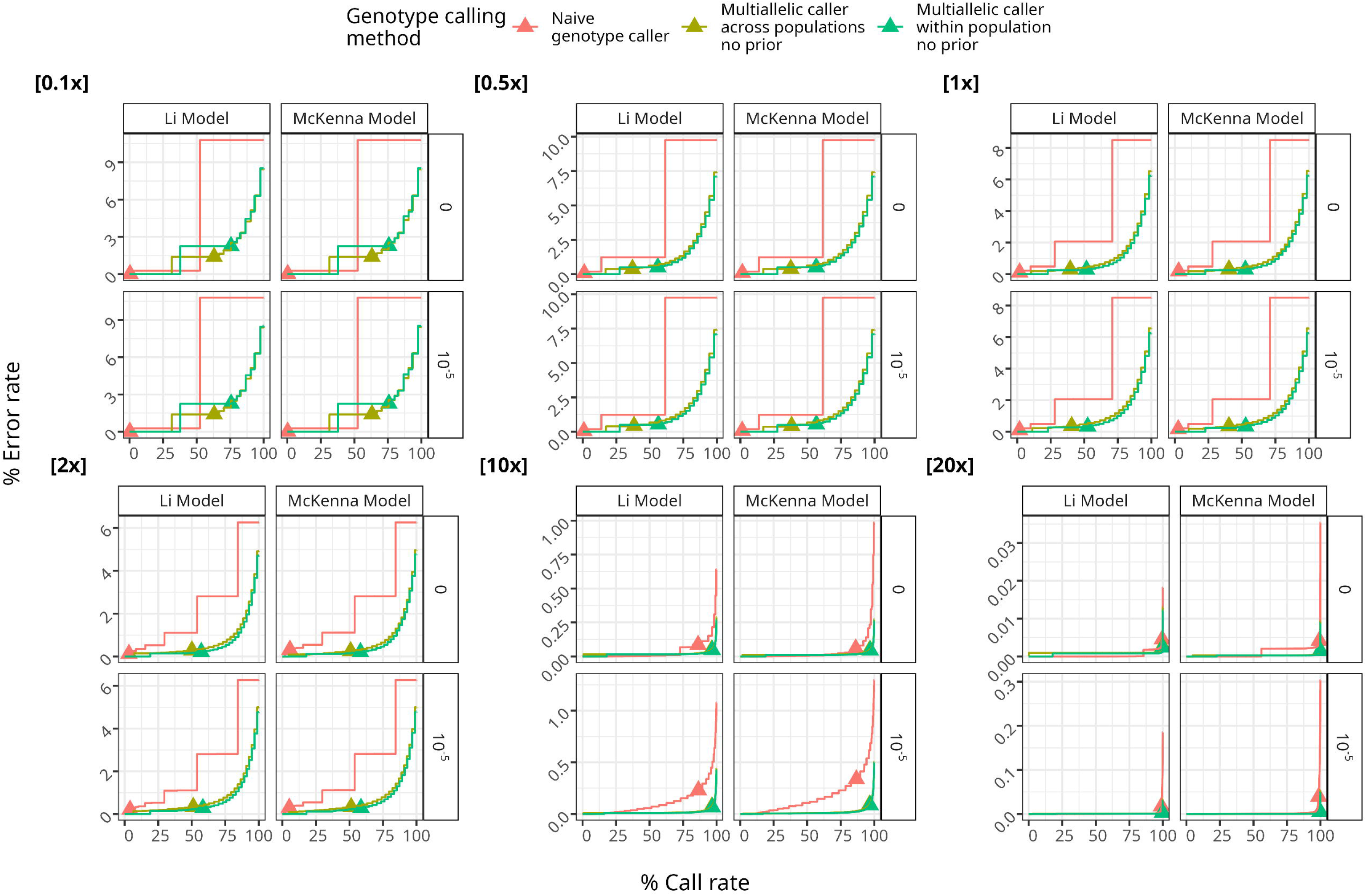

## References

Baumdicker, F. et al. (2022). Efficient ancestry and mutation simulation with msprime 1.0. Genetics, 220(3).

Caetano-Anolles, D. (2023). GVCF - Genomic Variant Call Format Technical Documentation.

da Fonseca, R. R. et al. (2016). Next-generation biology: Sequencing and data analysis approaches for non-model organisms. Mar. Genomics, 30, 3–13.

Danecek, P. et al. (2016). Multiallelic calling model in bcftools (-m). https://samtools.github.io/bcftools/call-m.pdf.

Danecek, P. et al. (2021). Twelve years of SAMtools and BCFtools. Gigascience, 10(2).

Fox, E. A. et al. (2019). ngsLD: evaluating linkage disequilibrium using genotype likelihoods. Bioinformatics, 35(19), 3855–3856.

Haller, B. C. and Messer, P. W. (2023). SLiM 4: Multispecies Eco-Evolutionary modeling. Am. Nat., 201(5), E127–E139.

Illumina (2014). GVCF files. https://support.illumina.com/help/BaseSpace_App_TumorNormal_help/Content/Vault/Informatics/Sequencing_Analysis/BS/swSEQ_mBS_gVCF.htm. Accessed: 2023-12-14.

Korneliussen, T. S. and Moltke, I. (2015). NgsRelate: a software tool for estimating pairwise relatedness from next-generation sequencing data. Bioinformatics, 31(24), 4009–4011.

Korneliussen, T. S. et al. (2014). ANGSD: Analysis of next generation sequencing data. BMC Bioinformatics, 15, 356.

Li, H. (2011). A statistical framework for SNP calling, mutation discovery, association mapping and population genetical parameter estimation from sequencing data. Bioinformatics, 27(21), 2987–2993.

Lou, R. N. et al. (2021). A beginner’s guide to low-coverage whole genome sequencing for population genomics. Mol. Ecol., 30(23), 5966–5993.

Luqman, H. et al. (2021). Identifying loci under selection via explicit demographic models.

Mas-Sandoval, A. et al. (2022). Fast and accurate estimation of multidimensional site frequency spectra from low-coverage highthroughput sequencing data. Gigascience, 11.

McKenna, A. et al. (2010). The genome analysis toolkit: a MapReduce framework for analyzing next-generation DNA sequencing data. Genome Res., 20(9), 1297–1303.

Meisner, J. and Albrechtsen, A. (2018). Inferring population structure and admixture proportions in Low-Depth NGS data. Genetics, 210(2), 719–731.

Messer, P. W. (2013). SLiM: simulating evolution with selection and linkage. Genetics, 194(4), 1037–1039.

Mölder, F. et al. (2021). Sustainable data analysis with snakemake. F1000Res., 10(33), 33.

Nielsen, R. et al. (2011). Genotype and SNP calling from next-generation sequencing data. Nat. Rev. Genet., 12(6), 443–451.

O’Rawe, J. A. et al. (2015). Accounting for uncertainty in DNA sequencing data. Trends Genet., 31(2), 61–66.

Rasmussen, M. S. et al. (2022). Estimation of site frequency spectra from low-coverage sequencing data using stochastic EM reduces overfitting, runtime, and memory usage.

Skotte, L. et al. (2013). Estimating individual admixture proportions from next generation sequencing data. Genetics, 195(3), 693–702.

Soraggi, S. et al. (2018). Powerful inference with the D-Statistic on Low-Coverage Whole-Genome data. G3, 8(2), 551–566.

Van der Auwera, G. A. and O’Connor, B. D. (2020). Genomics in the Cloud: Using Docker, GATK, and WDL in Terra. “O’Reilly Media, Inc.”.

Vieira, F. G. et al. (2016). Improving the estimation of genetic distances from Next-Generation sequencing data. Biol. J. Linn. Soc. Lond., 117(1), 139–149.

Wang, J. et al. (2016). Variation in linked selection and recombination drive genomic divergence during allopatric speciation of european and american aspens. Mol. Biol. Evol., 33(7), 1754–1767.

Waples, R. K. et al. (2019). Allele frequency-free inference of close familial relationships from genotypes or low-depth sequencing data. Mol. Ecol., 28(1), 35–48.

Zhao, D. et al. (2023). A genomic quantitative study on the contribution of the Ancestral-State bases relative to derived bases in the divergence and local adaptation of populus davidiana. Genes, 14(4).

Zhao, L. et al. (2022). distangsd: Fast and accurate inference of genetic distances for Next-Generation sequencing data. Mol. Biol. Evol., 39(6).

