## Supplementary Material for "vcfgl: A flexible genotype likelihood simulator for VCF/BCF files"

#### Supplemental content

**1 Section S1: Definitions** **4**

### Supplemental tables

1 Description of the fields in I16 tag . . . . . . . . . . . . . . . . . . . . . . 6

### Supplemental figures

1 . . . . . . . . . . . . . . . . . . . . . . . . . . . . . . . . . . . . . . . . . 17

1 Error rate versus call rate for different genotype calling methods (color) and GL models (line type) at different depths (rows) and beta distribution variance values (columns). . . . . . . . . . . . . . . . . . . . . . . . . . 18

2 Error rate versus call rate for different genotype calling methods, across read depths (panels), beta distribution variance values (facet rows), and GL models (facet columns). The genotype calling methods are: Naive genotype caller, BCFtools multiallelic caller across populations with no prior (`-P 0`), and BCFtools multiallelic caller within population (`-G`) with no prior (`-P 0`). The beta distribution variance values are 0 (precise quality scores) and $10^{-5}$ . . . . . . . . . . . . . . . . . . . . . . . . 18

3 Error rate versus call rate for different population pooling approaches (color) at different depths (panel), and beta distribution variance values (linetype) using the Li GL model. The two population pooling approaches are: BCFtools multiallelic caller across populations, and BCFtools multiallelic caller within populations using `-G` option. . . . . . . . . . . . . . 19

4 Error rate versus call rate for different population pooling approaches (color) at different depths (panel), and beta distribution variance values (linetype) using the McKenna GL model. The two population pooling approaches are: BCFtools multiallelic caller across populations, and BCFtools multiallelic caller within populations using `-G` option. . . . . 20

5 . . . . . . . . . . . . . . . . . . . . . . . . . . . . . . . . . . . . . . . . . 21

5 Error rate versus call rate for different genotype calling methods (columns) and read depths (rows). The vertical line segments below the 0 on the y axis denote the minimum GQ threshold of 20, and the horizontal line segments after the 100 on the x axis denote the final error rate. The colored solid lines denote the discordance rate for the four different types of discordance: HOM-HOM, HOM-HET, HET-HOM, and HET-HET, corresponding to discordant genotype calls where both the true and called genotypes are homozygous, where the true genotype is homozygous and the called genotype is heterozygous, where the true genotype is heterozygous and the called genotype is homozygous, and where both the true and called genotypes are heterozygous, respectively. . . . . . . . . 22
6 The proportion of variable sites retrieved by different genotype calling methods, as a function of sequencing depth, for different beta distribution variance values (color). The proportion of variable sites retrieved is calculated as the number of variable sites retrieved divided by the total number of truly variable sites in the simulated data, and multiplied by 100 to obtain a percentage. . . . . . . . . . . . . . . . . . . . . . . . . . . 23

### 1 Section S1: Definitions

#### 1.1 Phred-scaled quality scores

Phred-scaled quality score $Q_{Phred}$ represents the level of confidence associated with each base call in DNA sequencing and is logarithmically related to the sequencing error probabilities ($\mathbb{P}(e)$) (Ewing and Green, 1998).

$$Q_{Phred} = -10 \times log_{10}(\mathbb{P}(e)),$$

The resulting quality score, denoted as $Q_{Phred}$, is often capped by bioinformatics software, such as BCFtools (Danecek *et al.*, 2021). Secondly, they are also discretized and centralized for representation in text files: floor$(Q_{Phred}) + 0.499$. For instance, BCFtools uses a maximum quality score of 63 and a minimum quality score of 4 (BCFtools version 1.18, BCFtools/bam2bcf.c:381-382).

$$Q_{Phred} = \begin{cases} 4, & Q_{Phred} < 4 \\ 63, & Q_{Phred} > 63 \\ Q_{Phred}, & \text{otherwise.} \end{cases}$$

Where 63 corresponds to the probability of incorrect base call $\mathbb{P}$(e)= $5.011872 \cdot 10^{-07}$, thus a base call accuracy of 0.9999995, and 4 corresponds to $\mathbb{P}$(e)= 0.3981072 thus a base call accuracy of 0.601892.

A technical detail regarding the representation of quality scores is, the quality scores in FASTQ and SAM files use Phred+33 encoding, where 33 is added to the ASCII value of the raw score to obtain the resulting ASCII value.

#### 1.2 VCF tags

##### 1.2.1 QS Tag

The QS tag is an INFO tag containing the normalized phred-scale quality score sums. The elements in the QS tag correspond to the normalized quality score sums for the alleles as they appear in the REF and ALT fields at each genomic position. For each sample, the quality scores associated with the nucleotide bases in the REF and ALT fields are summed up to obtain a total quality score for that sample. This quality score sum $sum_s$ summarizes the overall sequencing quality at a specific genomic position within a sample. Then, the quality scores for each base in each sample are normalized by dividing the quality score of a base by the total quality score of the sample. This normalization aims to obtain the relative quality of each base, accounting for variations in sequencing quality across samples. Finally, these normalized scores are aggregated across all samples for each corresponding allele. This aggregation results in one quality score sum per allele, considering all available data, enabling a comprehensive overview of the quality scores.

For a genomic position with $A$ alleles and $S$ samples, the QS tag is calculated for each allele $a$ jointly for all samples as follows:

$$\text{sum}_s = \sum_{a=1}^{A} Q_{sa}$$

$$\text{norm}_{sa} = \frac{Q_{sa}}{\text{sum}_s}$$

$$\text{QS}[a] = \sum_{s=1}^{S} \text{norm}_{sa}.$$

##### 1.2.2 I16 Tag

The I16 tag is an INFO tag which is typically produced by BCFtools mpileup and necessary for genotype calling using BCFtools.

#### 1.3 Compatibility with other tools

The simulated output files from vcfgl is compatible with commonly-used tools such as BCFtools, GATK and ANGSD. The BCFtools compatibility was validated for

Table S 1: Description of the fields in I16 tag

| Index | Description |
|---|---|
| 1 | Number of reference bases on the forward strand |
| 2 | Number of reference bases on the reverse strand |
| 3 | Number of non-reference bases on the forward strand |
| 4 | Number of non-reference bases on the reverse strand |
| 6 | Sum of squares of reference base qualities |
| 7 | Sum of non-reference base qualities |
| 8 | Sum of squares of non-reference base qualities |
| 9 | Sum of reference mapping qualities |
| 10 | Sum of squares of reference mapping qualities |
| 11 | Sum of non-reference mapping qualities |
| 12 | Sum of squares of non-reference mapping qualities |
| 13 | Sum of tail distances for reference bases |
| 14 | Sum of squares of tail distances for reference bases |
| 15 | Sum of tail distances for non-reference bases |
| 16 | Sum of squares of tail distances for non-reference bases |

both `QS` and `I16` tags using `bcftools call -m` and `bcftools call -c`. The GATK compatibility was validated using the following command:

```
java -jar gatk-4.4.0.0/gatk-package-4.4.0.0-local.jar ValidateVariants
-validate-GVCF true -variant <INPUT> -validation-type-to-exclude REF.
```

#### 1.4 Genotype likelihood models

##### 1.4.1 McKenna model: Direct genotype likelihood model

The base-calling errors are assumed to be correct and independent in the McKenna diploid genotype likelihood model.

We can define genotype likelihood $P(D_{mn} \mid G)$ as the probability of the sequencing data observed in individual $n$ at site $m$, $D_{mn}$, conditional on a particular genotype $G_{mn}$. Given a per-site read depth of $X_{mn}$ representing the sequencing reads aligned to position $m$ for individual $n$, let $b_i$ be the nucleotide for the $i$th base read at site, and let $\epsilon_i$ be the sequencing error probability associated with the $i$th base.

$$
\begin{aligned}
\mathbb{P}(D_{mn} = b_i \mid G_{mn} = G) &= \prod_{i=1}^{X_{mn}} \mathbb{P}(b_i \mid G_{mn} = G), \\
&= \prod_{i=1}^{X_{mn}} \left( \frac{1}{2}\mathbb{P}(b_i \mid A_1) + \frac{1}{2}\mathbb{P}(b_i \mid A_2) \right).
\end{aligned}
$$

The probability of observed data given a genotype, namely the genotype likelihood, is therefore given by

$$
\mathbb{P}(b_i \mid G) = \begin{cases} \epsilon_i \times \frac{1}{3} & \text{if } b_i \neq A, \\ 1 - \epsilon_i & \text{if } b_i = A. \end{cases}
$$

##### 1.4.2 Li model: Genotype likelihood model with error correlation

Genotype likelihood model with correlated errors is described by Li *et al.* (2008) and is implemented in the SAMtools/BCFtools as follows:

Let $\mathbb{P}_l(m, n)$ be the probability of observing $m$ errors in $n$ reads, where

$$
\mathbb{P}_l(m, n) = \binom{n}{m} \times \epsilon_l^m \times (1 - \epsilon_l)^{n-m}
$$

and $\epsilon_l$ is the base-calling error of the read $i$ where the base call $b_i = l$.

Let us define function $R$ of the true genotype $G = \{g_1, g_2\}$ as

$$
R(G) = \begin{cases} \binom{n}{n_{g_i}} \times \frac{1}{2^n} & \text{if } g_1 \neq g_2 \\ 1 & \text{otherwise} \end{cases}
$$

where $n_{g_1}$ and $n_{g_2}$ is the number of reads supporting $g_1$ and $g_2$, respectively. In the case of heterozygous true genotype, the probabilities are approximated assuming that the errors are distributed between the nucleotides $g_1$ and $g_2$ with equal likelihood.

Then, the genotype likelihood can be expressed in terms of the read depths (i.e. the number of times a specific nucleotide is sequenced) assuming non-independent base calling error rates as

$$\mathbb{P}(b_i = l \mid G) = R(G) \times \prod_{i=1}^{n_l} \left( \frac{\sum_{m=i}^{n} \mathbb{P}_l(m, n)}{\mathbb{P}_l(0, n) + \sum_{m=i}^{n} \mathbb{P}_l(m, n)} \right)^{(\theta^{i-1} \times 0.97) + 0.03}$$

where the unknown parameter $\theta$ depends on a number of factors related to the sequencing and can be estimated from real data. In practice, $\theta = 0.85$ is a commonly used value as it was found as the best parameter value for Illumina Genetic Analyzer data (Li *et al.*, 2008). Our program allows the user to specify the theta parameter for their specific simulation setup.

### 2 Section S2: Simulation

#### 2.1 Simulating read depths

The Lander-Waterman theory argues that the number of reads covering a site can be approximated by a Poisson distribution under some statistical assumptions (Lander and Waterman, 1988). In this model, the reads are assumed to be uniformly sampled from the genome. Therefore, we sample the number of reads at each site from a Poisson distribution $X \sim \text{Pois}(\lambda)$

$$\mathbb{P}(X = x) = \frac{e^{-\lambda}\lambda^{x}}{x!}$$

where the $x$ is the number of reads covering a given site on the genome and $\lambda$ is the mean per-site read depth.

#### 2.2 Simulation at maximum read depth

Simulation of genotype likelihoods at highest possible depth values is typically used for comparing genotype calling-based methods with the genotype likelihood-based methods that are expected to be equivalent at maximum confidence. For this, we provide an option for the users to simulate genotype likelihoods at maximum confidence with little computational cost via `-depth inf` option.

represented as 8 bit unsigned integers (Bonfield *et al.*, 2021), which has a range of 0 to 255.

In absolute confidence simulations, we assign a genotype likelihood value of 0 to the true genotype and $-\infty$ to all other genotypes. Likewise, we set the genotype probability value of the true genotype to 1 and the probabilities of other genotypes to 0. Additionally, the phred scaled likelihood for the true genotype is set to 255, and 0 for other genotypes.

#### 2.3 Simulating the errors in the raw quality scores

We use the Beta distribution to simulate the errors in raw base quality scores. Beta distribution has a probability density function

$$f(x) = \begin{cases} \frac{\Gamma(\alpha+\beta)}{\Gamma(\alpha)\Gamma(\beta)} x^{\alpha-1}(1-x)^{\beta-1}, & \text{for } 0 \leq x \leq 1, \\ 0 & \text{otherwise,} \end{cases}$$

where $0 \leq x \leq 1$, $\alpha > 0$ and $\beta > 0$.

If $X \sim \text{Beta}(\alpha, \beta)$, then the mean of X, $\mu$, is given by

$$\mu = \text{E}[X] = \frac{\alpha}{\alpha + \beta}$$

and the variance of X, $\sigma^2$, is given by

$$\sigma^2 = \text{Var}(X) = \frac{\alpha\beta}{(\alpha+\beta)^2(\alpha+\beta+1)}.$$

We can directly re-parameterize the Beta distribution to get $\alpha$ and $\beta$ in terms of $\mu$ and $\sigma^2$,

$$\alpha = \mu \left( \frac{\mu(1-\mu)}{\sigma^2} - 1 \right)$$

$$\beta = (1-\mu) \left( \frac{\mu(1-\mu)}{\sigma^2} - 1 \right) = \alpha \left( \frac{1}{\mu} - 1 \right).$$

The parameters $\mu$ and $\sigma^2$ are defined by the user with `-error-rate` and `-beta-variance` options, respectively.

**Algorithm 1:** Simulation of genotype likelihoods with Beta-distributed base calling errors

**Input:** Genotypes $(A_1, A_2)$ at each site $s \in \{1, ..., S\}$ represented as combinations of ancestral (0) and derived (1) allelic states, $A_1, A_2 \in \{0, 1\}$. Average per-site read depth $d$. Base-calling error rate $e$. Beta distribution variance $v$.

**Output:** $L_s^{'}$ vector of genotype likelihoods at site $s$ as log10-transformed likelihood ratios.

$R_s \sim \text{Poisson}(d)$ ; `// sample number of reads to simulate (R)`

**if** $R_s = 0$ **then**

  $L_s^{'} \leftarrow$ '.' ; `// set site to missing`

**else**

  $\alpha = e\left(\frac{e(1-e)}{v} - 1\right), \beta = \alpha\left(\frac{1}{e} - 1\right)$ ;

  $\epsilon \sim \text{Beta}(\alpha, \beta)$ ; `// beta-distributed base calling quality`

  $$\mathbb{P}(b|A) = \begin{cases} \frac{\epsilon}{3}, & b \neq A \\ 1 - \epsilon, & \text{otherwise} \end{cases}$$

  $L_s(A_1, A_2) \leftarrow \log_{10}(\mathbb{P}(D|G = \{A_1, A_2\})) = \sum_{r=1}^{R} \log_{10}(\mathbb{P}(b_r|G = \{A_1, A_2\}))$ ;

  $L_s^{'} \leftarrow L_s - \max(L_s)$ ; `// normalize log likelihoods`

### 3 Section S3: Benchmarking

#### 3.1 Benchmarking pipeline

##### 3.1.1 Coalescence simulation

We used msprime and stdpopsim (Adrion *et al.*, 2020; Elise Lauterbur *et al.*, 2023; Baumdicker *et al.*, 2022; Kelleher *et al.*, 2016) to simulate the variable sites chromosome 22 with Discrete-Time Wright Fisher model (Nelson *et al.*, 2020) using a mutation rate of $2.35 \times 10^{-8}$, which in total resulted in $328,230$ variable sites. The simulated dataset consisted of a total of 100 diploid individuals, with 50 individuals from YRI and 50 individuals from CEU populations described in the Out of Africa model (Gutenkunst *et al.*, 2009). Watterson's theta estimates $\hat{\theta}$ for the simulated true data from msprime are calculated as 0.000982 for the YRI population (50 individuals), 0.000624 for the CEU population (50 individuals), and 0.001102 for the entire dataset (100 individuals).

##### 3.1.2 Genotype likelihood simulation

We used vcfgl to simulate the genotype likelihoods and errors in raw quality scores at varying read depths (20, 10, 2, 1, 0.5, 0.1), each with 20 simulation replicates. We used a mean error rate of 0.2%, along with Beta distribution variances of 0 (indicating no errors in quality scores), $1 \times 10^{-5}$, $1 \times 10^{-6}$, and $1 \times 10^{-7}$. The distributions corresponding to these variances are shown in figures **??**, **??**, and **??**, respectively.

#### 3.2 Genotype calling

##### 3.2.1 Naive genotype calling method

For the naive genotype calling method, we used the following command:

`bcftools +tag2tag - -GL-to-GT -threshold 1`

This command converts genotype likelihoods to genotype calls, where the genotype call is the genotype with the highest likelihood. The genotype probability threshold is set to 1 (i.e., no thresholds are applied). This is the simplest possible genotype calling method, and it is used as a baseline for comparison with other genotype calling methods.

##### 3.2.2 BCFtools multiallelic genotype caller

We used BCFtools multiallelic caller (Danecek *et al.*, 2016, 2021) to call genotypes with the default theta parameter value 0.0011 (which is described as an approximate typical value for humans (Danecek *et al.*, 2016)), and with the theta parameter disabled. We also compared two approaches: (1) genotype calling with all individuals pooled together, and (2) genotype calling per population. Overall, we used the following four commands:

`bcftools call -m -prior 0.0011` // across populations

`bcftools call -m -prior 0` // across populations

`bcftools call -m -prior 0.0011 -G {popinds_list}` // within populations

`bcftools call -m -prior 0 -G {popinds_list}` // within populations

#### 3.3 Quantification of the genotype calling accuracy

To evaluate the genotype calling accuracy, we defined two metrics, error rate ($E$) and call rate ($C$), based on the number of discordant and concordant genotype calls between the true genotypes and the genotype calls at different genotype quality (GQ) thresholds.

For this, we calculated the number of discordant and concordant genotype calls for each sample with a given GQ value. Then, the resulting number of discordant and concordant calls were summed across all samples, and the mean of the results across all replicates was used to obtain the average discordance and concordance rates at each GQ bin. The calculation of these basic metrics is implemented in an efficient utility program named `gtDiscordance` and is freely available as a utility tool in vcfgl.

##### 3.3.1 Genotype qualities (GQ)

For the naive genotype calling method, the second smallest phred-scaled likelihood (PL) value was assigned as the GQ value, where the PL values are assumed to be rescaled so that the smallest PL value is 0.

For the sites identified as invariable by the BCFtools multiallelic caller, we assigned the highest GQ value as the GQ value for non-SNP sites is not reported by BCFtools.

##### 3.3.2 Error rate $E$ and call rate $C$

The error rate $E$ was calculated cumulatively over the data points sorted by GQ in descending order. The cumulative error rate $E$ was calculated as the cumulative number of discordant genotype calls up to the current GQ bin, divided by the total number of genotype calls made by the genotype caller. This value is referred to as the error rate in the figures. The call rate $C$ was calculated as the cumulative number of genotype calls up to the current GQ bin, divided by the total number of genotype calls that were made by the genotype caller. This value is referred as the call rate in the figures.

Let $N_D$ be the number of discordant sites, defined as:

$$N_D = \sum_{s \in G_C} I(s)$$

where $I(s) = 1$ if $G_{C,s} \neq G_{T,C,s}$ and $I(s) = 0$ otherwise.

Let $N_C$ be the number of concordant sites, defined as:

$$N_C = \sum_{s \in G_C} (1 - I(s)).$$

Given a dataset grouped by depth, genotype likelihood model, beta variance, genotype calling method, the error rate $E$ and the call rate $C$ were calculated cumulatively for each GQ bin, indexed by $Q$, as $E_Q$ and $C_Q$, where the GQ bins are arranged in descending order, $GQ = \{1, \ldots, 129\}$.

$$C_Q = \frac{\sum_{i=1}^{Q} N_{D,i} + \sum_{i=1}^{Q} N_{C,i}}{\sum_{i=1}^{Q} N_{D,i} + \sum_{i=1}^{Q} N_{C,i}},$$

$$E_Q = \frac{\sum_{i=1}^{Q} N_{D,i}}{\sum_{i=1}^{Q} N_{D,i} + \sum_{i=1}^{Q} N_{C,i}}.$$

where $N_{D,i}$ and $N_{C,i}$ are the cumulative numbers of discordant and concordant sites, respectively, up to the GQ bin $i$.

#### 3.4 Results

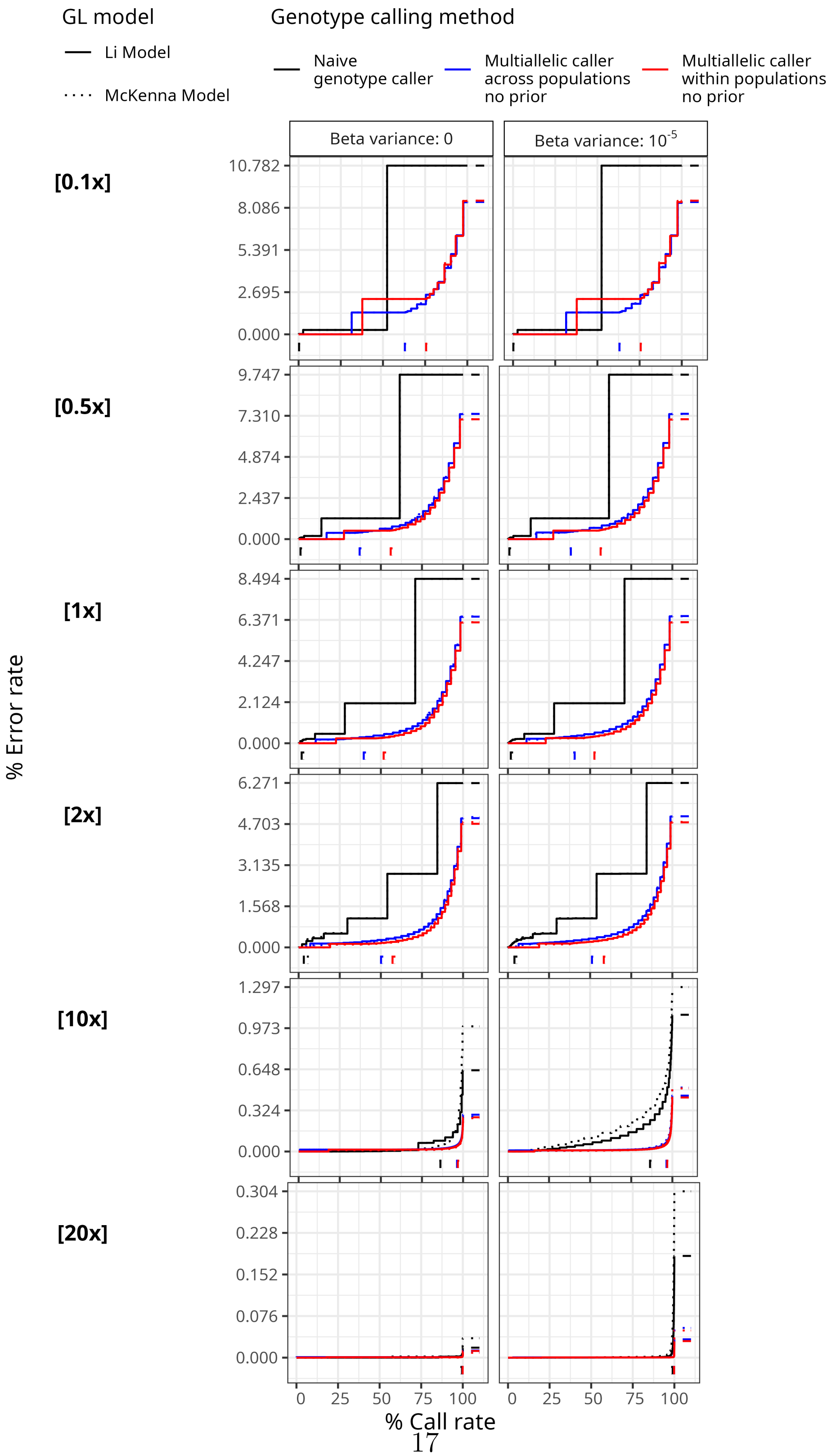

GL model
Li Model
McKenna Model
Genotype calling method
Naive genotype caller
Multiallelic caller across populations no prior
Multiallelic caller within populations no prior
Beta variance: 0
Beta variance: 10^-5
[0.1x]
[0.5x]
[1x]
[2x]
[10x]
[20x]
% Error rate
% Call rate

Figure S 1 *(previous page)*: Error rate versus call rate for different genotype calling methods (color) and GL models (line type) at different depths (rows) and beta distribution variance values (columns).

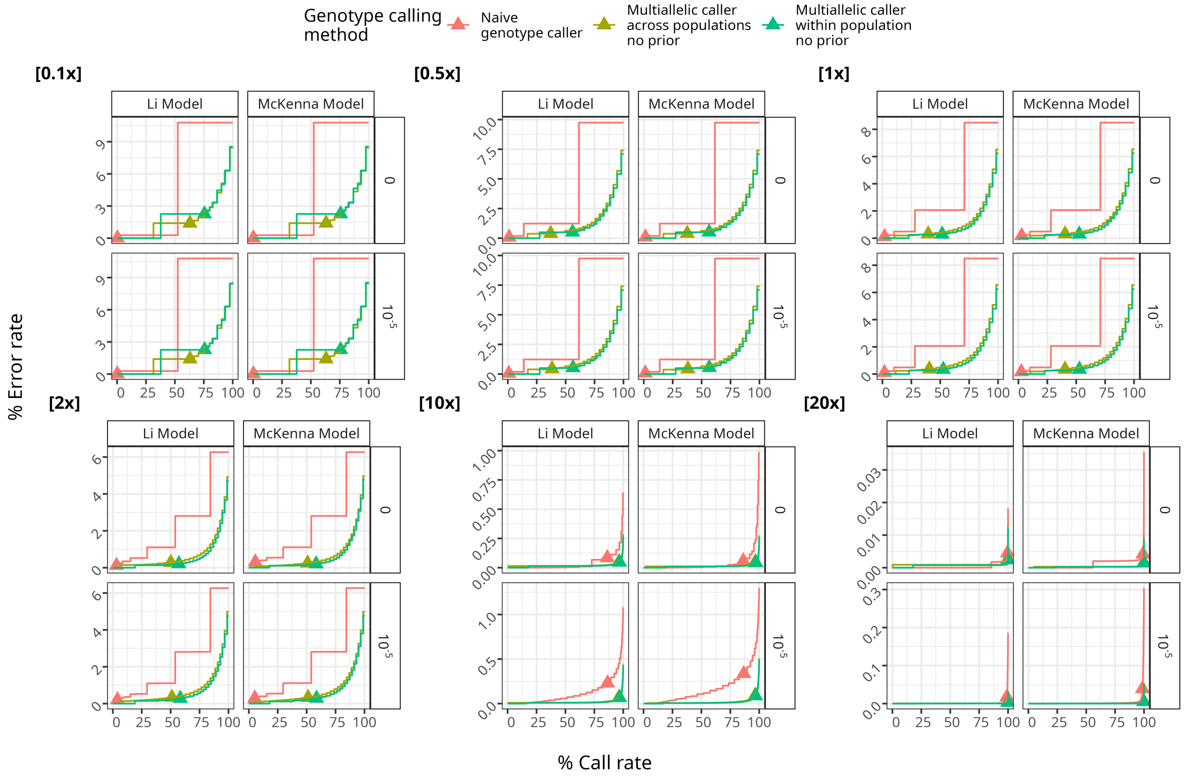

Figure S 2: Error rate versus call rate for different genotype calling methods, across read depths (panels), beta distribution variance values (facet rows), and GL models (facet columns). The genotype calling methods are: Naive genotype caller, BCFtools multiallelic caller across populations with no prior (`-P 0`), and BCFtools multiallelic caller within population (`-G`) with no prior (`-P 0`). The beta distribution variance values are 0 (precise quality scores) and $10^{-5}$

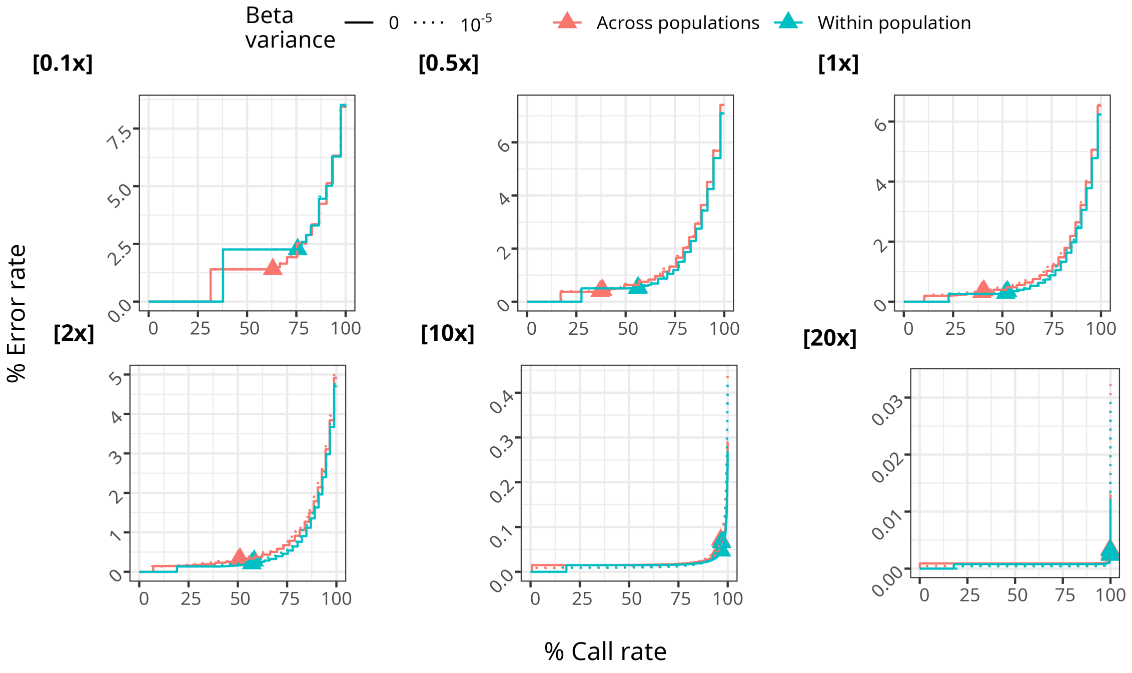

Figure S 3: Error rate versus call rate for different population pooling approaches (color) at different depths (panel), and beta distribution variance values (linetype) using the Li GL model. The two population pooling approaches are: BCFtools multiallelic caller across populations, and BCFtools multiallelic caller within populations using `-G` option.

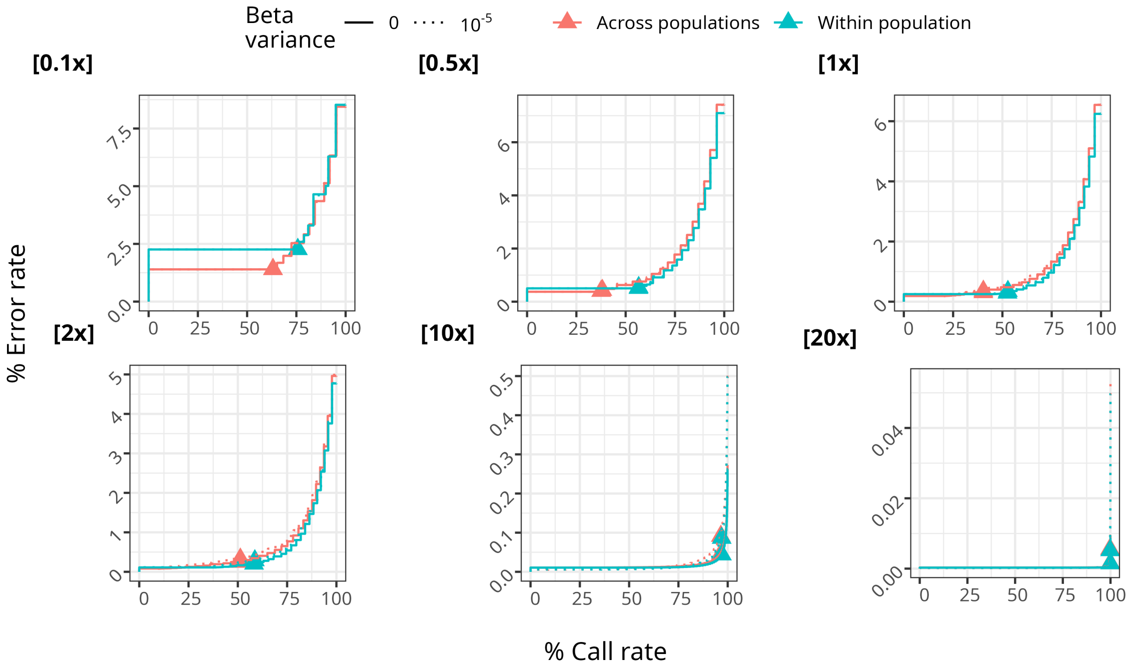

Figure S 4: Error rate versus call rate for different population pooling approaches (color) at different depths (panel), and beta distribution variance values (linetype) using the McKenna GL model. The two population pooling approaches are: BCFtools multiallelic caller across populations, and BCFtools multiallelic caller within populations using `-G` option.

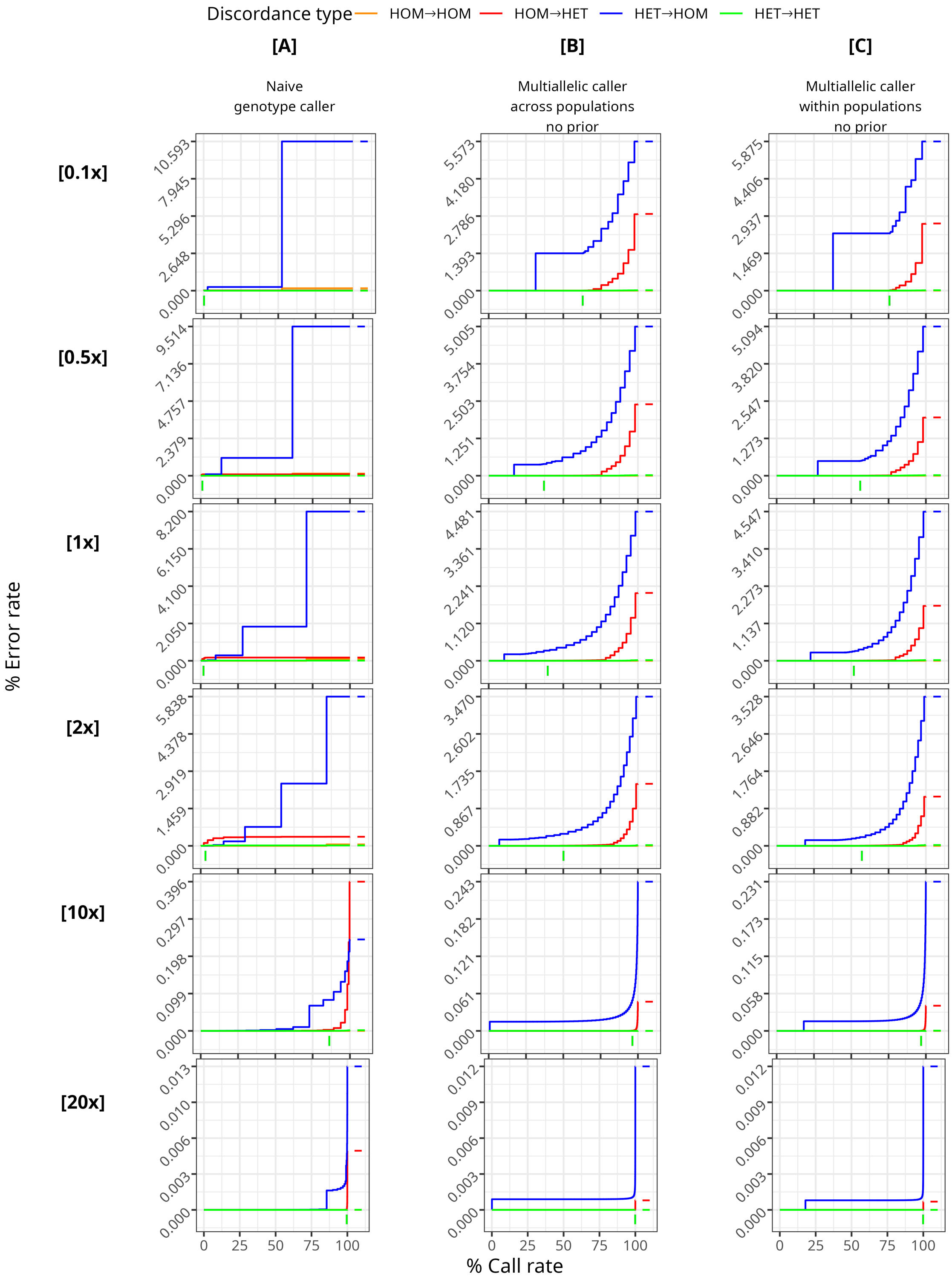
Discordance type
HOM→HOM
HOM→HET
HET→HOM
HET→HET
[A]
Naive genotype caller
[B]
Multiallelic caller across populations no prior
[C]
Multiallelic caller within populations no prior
[0.1x]
[0.5x]
[1x]
[2x]
[10x]
[20x]
% Error rate
% Call rate
0
25
50
75
100

Figure S 5 *(previous page)*: Error rate versus call rate for different genotype calling methods (columns) and read depths (rows). The vertical line segments below the 0 on the y axis denote the minimum GQ threshold of 20, and the horizontal line segments after the 100 on the x axis denote the final error rate. The colored solid lines denote the discordance rate for the four different types of discordance: HOM-HOM, HOM-HET, HET-HOM, and HET-HET, corresponding to discordant genotype calls where both the true and called genotypes are homozygous, where the true genotype is homozygous and the called genotype is heterozygous, where the true genotype is heterozygous and the called genotype is homozygous, and where both the true and called genotypes are heterozygous, respectively.

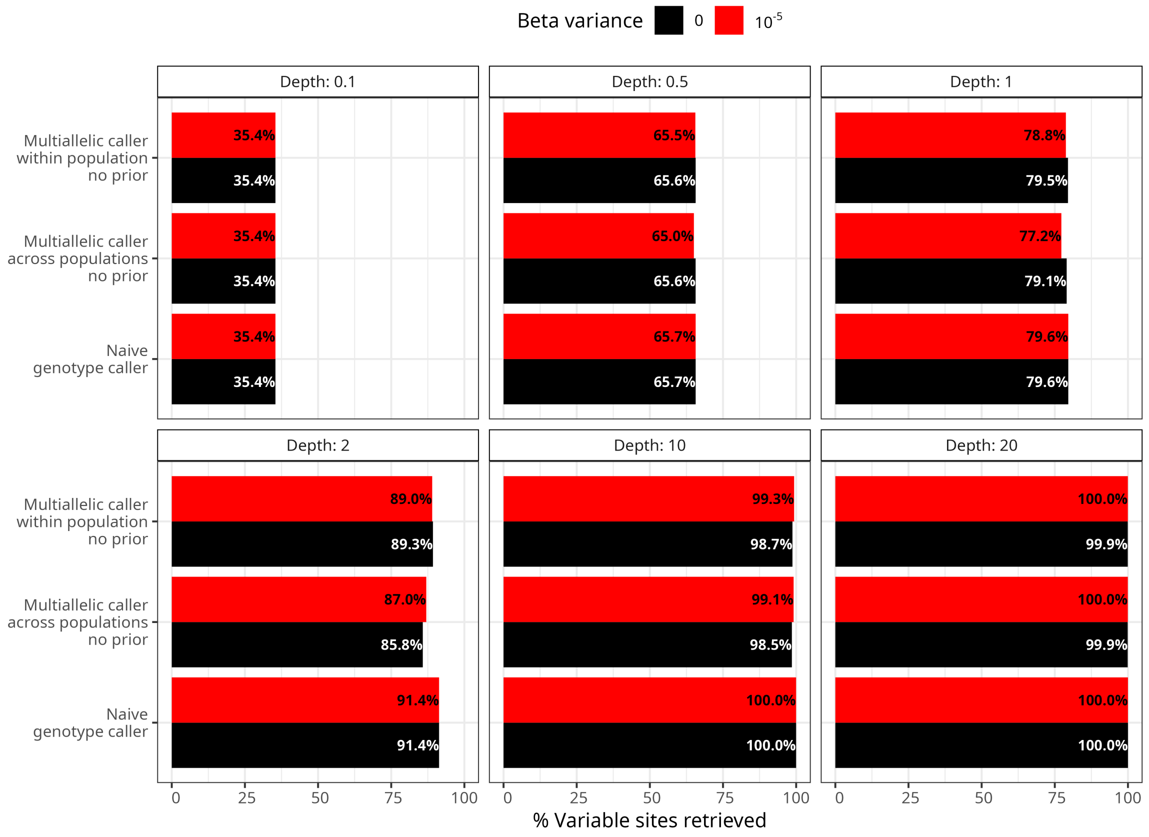

Figure S 6: The proportion of variable sites retrieved by different genotype calling methods, as a function of sequencing depth, for different beta distribution variance values (color). The proportion of variable sites retrieved is calculated as the number of variable sites retrieved divided by the total number of truly variable sites in the simulated data, and multiplied by 100 to obtain a percentage.

#### 3.5 Reproducibility and availability

We developed the benchmarking pipeline using Snakemake (Mölder *et al.*, 2021) to ensure reproducibility. The pipeline is freely available at https://github.com/isinaltinkaya/vcfgl_benchmarking.

### References

Adrion, J. R. *et al.* (2020). A community-maintained standard library of population genetic models. *Elife*, **9**.

Baumdicker, F. *et al.* (2022). Efficient ancestry and mutation simulation with msprime 1.0. *Genetics*, **220**(3).

Bonfield, J. K. *et al.* (2021). HTSlib: C library for reading/writing high-throughput sequencing data. *Gigascience*, **10**(2).

Danecek, P. *et al.* (2016). Multiallelic calling model in bcftools (-m). https://samtools.github.io/bcftools/call-m.pdf.

Danecek, P. *et al.* (2021). Twelve years of SAMtools and BCFtools. *Gigascience*, **10**(2).

Elise Lauterbur, M. *et al.* (2023). Expanding the stdpopsim species catalog, and lessons learned for realistic genome simulations.

Ewing, B. and Green, P. (1998). Base-calling of automated sequencer traces using phred. II. error probabilities. *Genome Res.*, **8**(3), 186–194.

Gutenkunst, R. N. *et al.* (2009). Inferring the joint demographic history of multiple populations from multidimensional SNP frequency data. *PLoS Genet.*, **5**(10), e1000695.

Kelleher, J. *et al.* (2016). Efficient coalescent simulation and genealogical analysis for large sample sizes. *PLoS Comput. Biol.*, **12**(5), e1004842.

Lander, E. S. and Waterman, M. S. (1988). Genomic mapping by fingerprinting random clones: a mathematical analysis. *Genomics*, **2**(3), 231–239.

Li, H. *et al.* (2008). Mapping short DNA sequencing reads and calling variants using mapping quality scores. *Genome Res.*, **18**(11), 1851–1858.

Mölder, F. *et al.* (2021). Sustainable data analysis with snakemake. *F1000Res.*, **10**, 33.

Nelson, D. *et al.* (2020). Accounting for long-range correlations in genome-wide simulations of large cohorts. *PLoS Genet.*, **16**(5), e1008619.
